# A synthetic biology approach to transgene expression

**DOI:** 10.1101/2023.08.31.555539

**Authors:** Philip T. Leftwich, Jessica C. Purcell, Michelle A.E. Anderson, Rennos Fragkoudis, Sanjay Basu, Gareth Lycett, Luke Alphey

## Abstract

The ability to control gene expression is pivotal in genetic engineering and synthetic biology. However, in most non-model and pest insect species, empirical evidence for predictable modulation of gene expression levels is lacking. This knowledge gap is critical for genetic control systems, particularly in mosquitoes, where transgenic methods offer novel routes for pest control. Commonly, the choice of RNA polymerase II promoter (Pol II) is the primary method for controlling gene expression, but the options are limited.

To address this, we developed a systematic approach to characterize modifications in translation initiation sequences (TIS) and 3’ untranslated regions (UTR) of transgenes, enabling the creation of a toolbox for gene expression modulation in mosquitoes and potentially other insects. The approach demonstrated highly predictable gene expression changes across various cell lines and promoter sequences, representing a significant advancement in mosquito synthetic biology gene expression. tools.

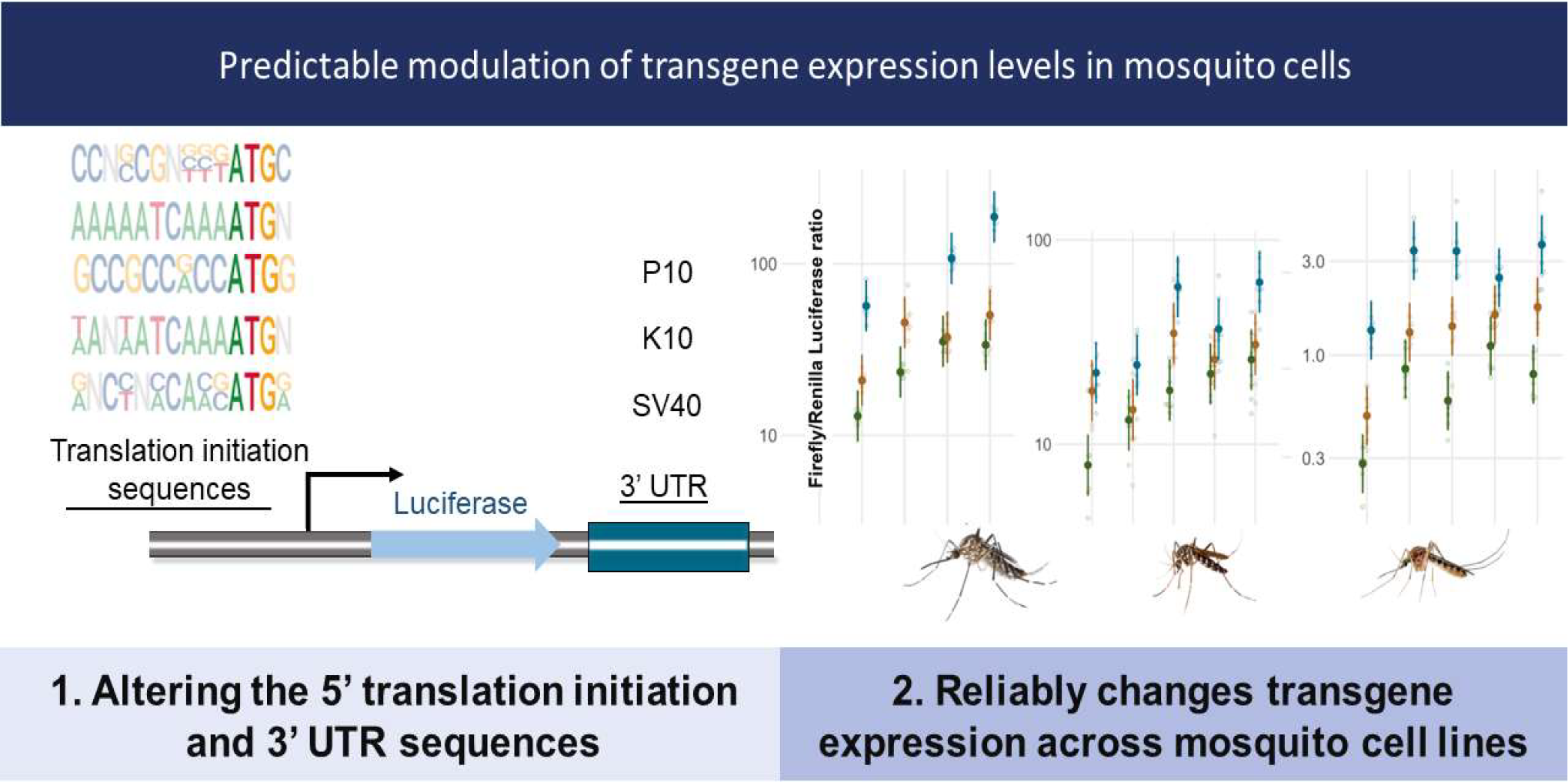

## Introduction

The ability to control the strength of expression of transgenes in a species of interest has underpinned genetic engineering and synthetic biology from their conception. There is, however, a lack of robust, empirical evidence for predictable modulation of gene expression levels in most non-model and pest insect species ^1,2^. In mosquitoes, transgenic methods afford novel routes for pest control however, genetic control systems depend on precise gene expression, so this lack of information is a critical gap in our technical capability. Commonly the selection of RNA polymerase II promoter (Pol II) can dictate the amount, timing, and spatial specificity of gene transcription. A limited number of promoters of viral origin are sometimes used, as these are active across a range of insect species, however there are applications for characterising endogenous promoters of varying activity. The choice of promoter is often determined by a requirement for specific spatial and/or temporal regulation, with few options for controlling expression level by this route beyond bespoke analysis of endogenous promoters.

Other methods commonly employed for modulating gene expression involve modifications to the mRNA sequences of the 5’ and 3’ untranslated regions (UTR), flanking the coding sequence of a transgene. Although these regions do not contribute to the final protein, they play crucial roles in mRNA stability and translation efficiency. In particular, the translation initiation sequence (TIS), a short (∼10nt) segment within the 5’ UTR just upstream of the start codon, has been shown to significantly impact translation efficiency in both vertebrates and invertebrates^3-6^. By using different TIS sequences, predictable changes in transgene expression can be achieved^2, 6, 7^.

The 3’UTR is more closely associated with mediating the termination of translation and ensuring efficient recycling of the translation complex, enabling multiple translations from the same mRNA molecule^8, 9^. In insect transgenesis, exogenous 3’UTR sequences are routinely used, including the viral-derived simian virus 40 (SV40) 3’UTR^11^, the P10 baculovirus 3’ UTR from *Autographa californica nucleopolyhedrovirus* (AcNPV)^12^ and the 3’UTR of the *K10* gene from *Drosophila melanogaster*^*13*^.

Despite the widespread use of these 3’UTR sequences, their relative efficacies are often based on anecdotal evidence. Strategically manipulating these untranslated regions provides a valuable approach to finely tune and optimize transgene expression in insect transgenesis.

We developed a systematic approach to characterising TIS and 3’UTR modifications to transgenes to build a toolbox for modulating gene expression in mosquitoes, and potentially other insects, when combined with viral or endogenous culicine promoter sequences. This toolbox provides an efficient way of expanding available Pol II promoters and affords routes to generating better regulation of activity of promoters across the species barrier. We initially tested the activities of the viral promoter HR5-IE1^10^ with a fully factorial combination of TIS and 3’UTR sequences in *Aedes aegypti, Aedes albopictus, Culex quinquefasciatus* and *Spodoptera frugiperda* cell lines. We then developed this further by taking the TIS/3’UTR combinations that produced the highest, lowest and median expression levels and demonstrated the highly replicable gene expression modulation effects in a panel of promoters.

We found that gene expression changes are highly predictable across a wide range of cell lines and promoter sequences. In conjunction with the characterisation of several endogenous culicine promoters, this represents a significant advance in the available gene expression tools for mosquito synthetic biology.

## Results and Discussion

We first measured the activity of five translation initiation sequences (TIS) (Table S1; BmLo, Syn21, Kozak, BmHi and Lep)^1–3,15^ and three different 3’UTRs (K10, SV40, P10)^11-13^ in a fully factorial design all downstream of the HR5-IE1 promoter. In total, we produced 15 different constructs and tested these in five different insect cell lines, from three disease relevant Culicine mosquito species (*A. aegypti (Aag2), A. albopictus (U4*.*4 & C6*.*36)* and *C. quinquefasciatus (Hsu)*) and one Lepidopteran species (*S. frugiperda* (*Sf9*)).

We found a highly replicable pattern of gene expression modulation across all five tested cell lines (Figure 1). Averaged across cell lines, the choice of TIS could produce up to a 2.55 relative-fold change in luciferase expression (95% CI 2.28-2.84; Table S2), while the choice of 3’UTR produced up to a 4.88 relative-fold change in luciferase expression (95% CI 4.52-5.26; Table S2).

**Figure 1.**
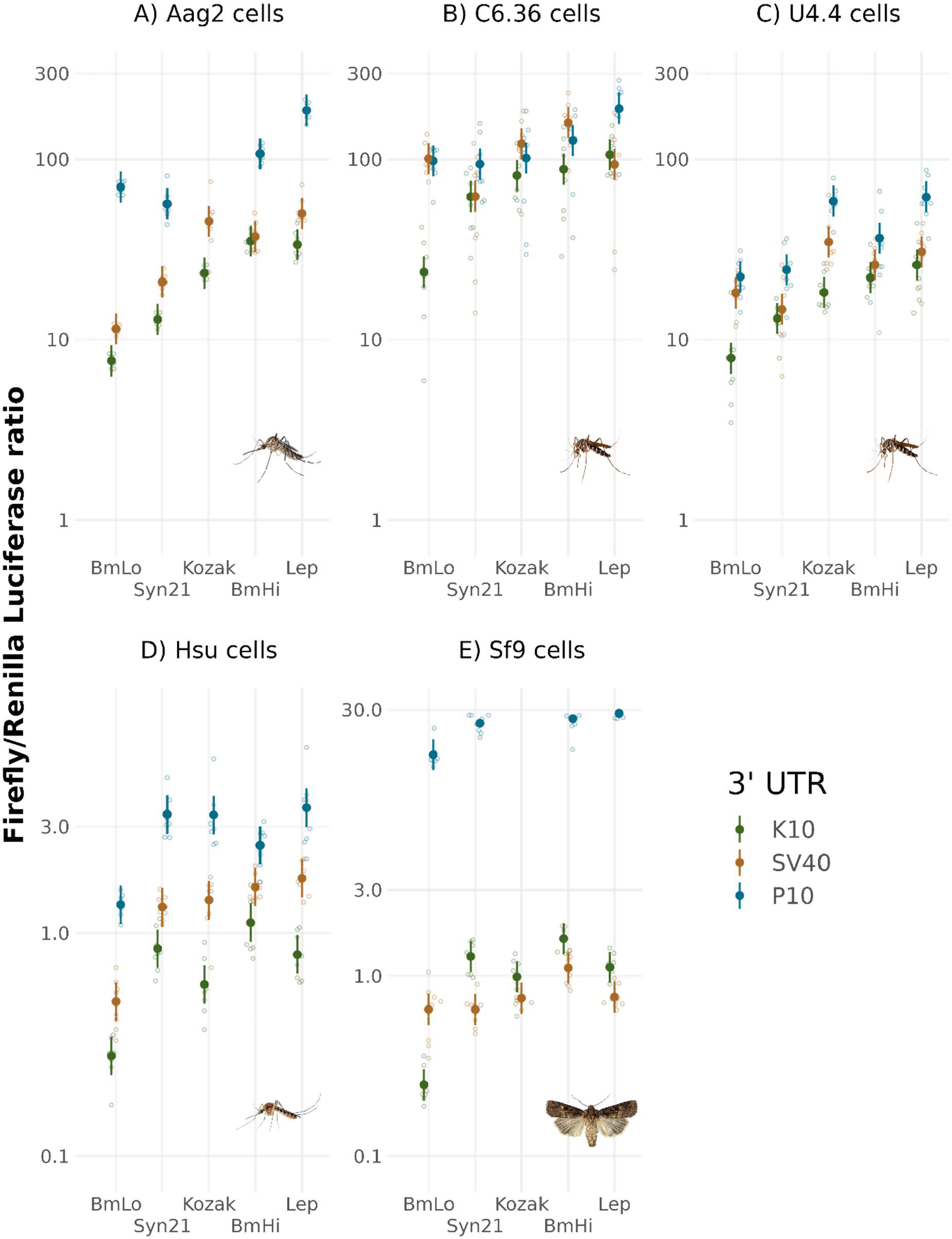
Combinations of translation initiation sequences (TIS) and 3’UTRs produce highly replicable gene expression across a range of insect cell lines. Ratios of FF luciferase compared to a RL control were used to measure activity, UTRs are organised left to right by average relative activity and nested within TIS also organised left to right by average relative activity. Large symbols and error bars represent mean and associated approximate 95% confidence intervals estimated with a generalized linear mixed model with a Gamma error distribution, raw data is shown as small open circles.

The estimates from our analysis indicate that TIS sequences are mainly insensitive to cell type and behave remarkably consistently (F_16,518_ = 11, P <0.001; Table S3). By contrast, the effect of the 3’UTR sequence on transgene expression was much more strongly affected by cell type (F_8,518_ = 250, P <0.001; Table S3). Expression from constructs with P10 had notably higher expression than expected in Sf9 cells and much lower in both C6/36 and U4.4 cells. Significant interactions between TIS sequences and 3’UTR sequences were small (F_8,518_ = 20, P <0.001; Table S3) therefore transcriptional activity appears to be primarily an additive effect when pairing TIS and 3’UTR sequences. This makes the “plug and play” notion of pairing different synthetic components together highly attractive, as effects on transgene expression are highly predictable.

The TIS/3’UTR combinations with the lowest (BmLo-K10) and highest (Lep-P10) expressions (18.2 (95% CI 13-22) relative-fold expression difference); were the same across all cell lines (Figure 1). To generalise our results, we decided to expand the range of promoter sequences tested by taking the BmLo-K10, Lep-P10 and Kozak-SV40 combinations and testing them with additional promoter sequences. In total, we tested seven regulatory sequences from four endogenous promoters from Culicine mosquitoes: two variants of the *hsp83* promoter^16^ (1.4kb & 888bp) (AAEL011708), from *A. aegypti* with the large (c.4.2kb) intronic sequence of the 5’UTR truncated to retain minimal acceptor and donor regions, three engineered variants of the *Polyubiquitin* promoter (AAEL003888) from *A. aegypti*^17^, along with *Polyubiquitin* from *C. quinquefasciatus* (CPIJ010919) and we demonstrate the first use case for a new endogenous promoter *A. albopictus* derived *Polyubiquitin* (AALF002118). These, along with OpIE2 were tested in Aag2 and U4.4 cells.

As expected, promoters of viral origin behaved very consistently across both cell lines, while endogenous promoters responded in a more cell-specific manner (Figure 2, Table S4). The *hsp83* promoter sequences produced equivalent levels of gene expression to OpIE2 when in Aag2 cells, but this was lower in U4.4 cells. Shortening this sequence by removing c.500bp upstream produced no significant reduction in gene expression. Polyubiquitin-derived promoter sequences generally produced the highest levels of gene expression; fluctuations in the strength of expression across cell lines may reflect the evolutionary origins of each sequence (*C. quinquefasciatus* derived sequence displayed lower activity than *A. albopictus* for example).

**Figure 2.**
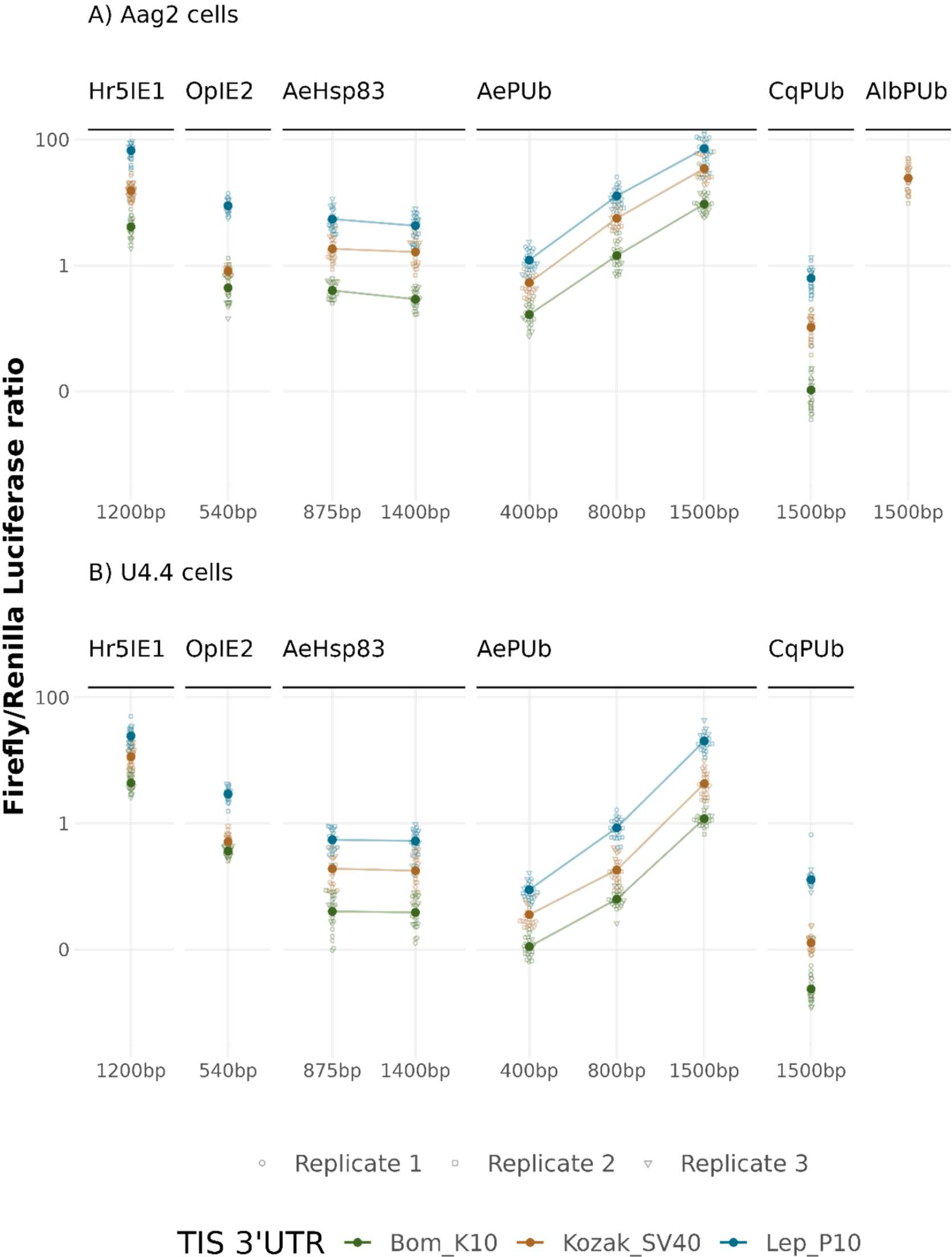
Fine-scale modulation of transgene expression with TIS/3’UTR combinations is highly replicable across a range of synthetic and endogenous promoters in two Culicine mosquito cell lines A) Aag2 (A. aegypti) cells B) A. albopictus-derived U4.4 cells. Ratios of FF luciferase compared to a RL control were used to measure activity, TIS/3’UTR combinations are organised left to right by average relative activity. Large solid symbols and error bars represent mean and associated approximate 95% confidence intervals estimated with a generalized linear mixed model with a Gamma error distribution; raw data is shown as open symbols. Lines connect promoters of the same origin.

The effects of TIS/3’UTR modification on gene expression were remarkably consistent for all promoter sequences. We found minimal differences in the responses between different regulatory sequences and TIS/3’UTR combinations (Promoter: F_8,1207_ = 354.91, P <0.001; TIS/3’UTR: F_2,1207_ = 509.96, P <0.001; Interaction effect: F_14,1193_ = 7.26, P <0.001; Figure 2, Table S5), indicating that these act largely independently.

We have demonstrated a straightforward method for modulating transgene expression in Culicine mosquitoes in a combinatorial approach that allows fine-scale manipulation. In these experiments, we have shown that TIS and 3’UTR sequences have highly predictable outcomes on transgene expression regardless of promoter sequence or cell line. These findings will likely translate well to whole-organism transgene expression^1^, and we anticipate this will provide a valuable resource to those working in synthetic biology, genetic modification, and mosquito genetic control.

## Methods

### Plasmids, cells, transfections and luciferase assay

Cells were seeded in 96-well plates one day before transfection with TransIT-PRO transfection kit (Mirus Bio, Madison, WI, US) according to manufacturer’s recommendations. Master mixes were prepared for eight wells of a 96-well plate, as replicate wells per experimental construct. This was repeated in three replicate experiments. Per well, transfection amounts are listed for each cell line in Supporting Information. Complete plasmid sequences are currently available on Github and will be available through NCBI.

Two days after transfection, cells were washed twice with ion-free PBS, lysed with 1X Passive Lysis Buffer then analysed using the Dual-Luciferase Assay kit on a GloMax multi+ plate reader (Promega, Southampton, UK).

General cell maintenance and plasmid information is described in Supporting Information (Tables S6-S9).

### Data Analysis

Scripts and raw data can be found at Github (https://github.com/Philip-Leftwich/Pol2-promoters). Complete information on analyses can be found in Supporting Information.

## Acknowledgements

This research was funded in part by research grants from the Wellcome Trust [110117/Z/15/Z and 226721/Z/22/Z], and the Bill & Melinda Gates Foundation [INV-008549] to LA; and by the Defense Advanced Research Projects Agency (DARPA) Safe Genes Program [N66001-17-2-4054] to Kevin Esvelt at MIT. JP was supported by a studentship from The Pirbright Institute (BBS/E/I/00001985). LA was also supported through strategic funding from the UK Biotechnology and Biological Sciences Research Council (BBSRC) to The Pirbright Institute (BBS/E/I/00007033, BBS/E/I/00007038 and BBS/E/I/00007039).For the purpose of open access, the author has applied a CC BY public copyright licence to any Author Accepted Manuscript version arising from this submission. The findings and conclusions contained within are those of the authors and do not necessarily reflect positions or policies of the Bill & Melinda Gates Foundation. The views, opinions and/or findings expressed are those of the authors and should not be interpreted as representing the official views or policies of the Department of Defense or the U.S. Government. The funders had no role in study design, data collection and analysis, decision to publish, or preparation of the manuscript

## Notes

https://github.com/Philip-Leftwich/Pol2-promoters

### Competing Interest Statement

The authors have declared no competing interest.

