## Supplemental methods for "A synthetic biology approach to transgene expression"

### Analysis

We carried out all analyses in R version 4.3.1 {R Development Core Team)

Luciferase readings were normalized for transfection by dividing the firefly activity by the Renilla activity. Data were analyzed by generalized linear models using a gamma error distribution with a log link, we included promoter, translation initation sequence, 3’UTR and cell line as factorial explanatory variables with all possible interactions. There is minor overdispersion because of heterogeneity among cell types, likely caused by the different basal expression levels of the OpIE2 promoter expressing Renilla luciferase. After each model was fitted, we used the ‘emmeans’ package (Lengh 2023) to back transform regression coefficients and calculate the estimated normalised mean expression with approximate 95% confidence intervals for each construct.

##### Experiment One

**Table S1. Fixed effects table for the generalized linear model (GLMM) fitted to the Luciferase Ratio detected in the engineered plasmids in experiment one – testing Hr5IE1 promoter with a combination of TIS, 3’UTR and cell types**

|  | **Value** | | |
| --- | --- | --- | --- |
| *Predictors* | *Estimates* | *CI* | *p* |
| (Intercept) | 35.34 | 29.18 – 43.36 | **<0.001** |
| Context [BmLo] | 0.22 | 0.16 – 0.29 | **<0.001** |
| Context [Kozak] | 0.66 | 0.50 – 0.88 | **0.004** |
| Context [Lep] | 0.95 | 0.72 – 1.26 | 0.745 |
| Context [Syn21] | 0.37 | 0.28 – 0.49 | **<0.001** |
| UTR [P10] | 3.05 | 2.30 – 4.03 | **<0.001** |
| UTR [SV40] | 1.06 | 0.80 – 1.40 | 0.695 |
| Cell Line [C6.36..Ae..albopictus.] | 2.50 | 1.89 – 3.31 | **<0.001** |
| Cell Line [Hsu..Cu..quinquefasciatus.] | 0.03 | 0.02 – 0.04 | **<0.001** |
| Cell Line [Sf9..S..frugiperda.] | 0.05 | 0.03 – 0.06 | **<0.001** |
| Cell Line [U4.4..Ae..albopictus.] | 0.63 | 0.47 – 0.83 | **0.001** |
| Context [BmLo] × UTR [P10] | 3.02 | 2.03 – 4.49 | **<0.001** |
| Context [Kozak] × UTR [P10] | 1.94 | 1.30 – 2.88 | **0.001** |
| Context [Lep] × UTR [P10] | 1.83 | 1.23 – 2.72 | **0.003** |
| Context [Syn21] × UTR [P10] | 1.44 | 0.97 – 2.14 | 0.071 |
| Context [BmLo] × UTR [SV40] | 1.42 | 0.96 – 2.11 | 0.082 |
| Context [Kozak] × UTR [SV40] | 1.84 | 1.24 – 2.73 | **0.003** |
| Context [Lep] × UTR [SV40] | 1.41 | 0.95 – 2.10 | 0.089 |
| Context [Syn21] × UTR [SV40] | 1.53 | 1.03 – 2.27 | **0.036** |
| Context [BmLo] × Cell Line [C6.36..Ae..albopictus.] | 1.24 | 0.84 – 1.85 | 0.283 |
| Context [Kozak] × Cell Line [C6.36..Ae..albopictus.] | 1.39 | 0.94 – 2.07 | 0.102 |
| Context [Lep] × Cell Line [C6.36..Ae..albopictus.] | 1.26 | 0.85 – 1.87 | 0.250 |
| Context [Syn21] × Cell Line [C6.36..Ae..albopictus.] | 1.92 | 1.29 – 2.86 | **0.001** |
| Context [BmLo] × Cell Line [Hsu..Cu..quinquefasciatus.] | 1.17 | 0.79 – 1.73 | 0.445 |
| Context [Kozak] × Cell Line [Hsu..Cu..quinquefasciatus.] | 0.79 | 0.53 – 1.18 | 0.255 |
| Context [Lep] × Cell Line [Hsu..Cu..quinquefasciatus.] | 0.75 | 0.51 – 1.12 | 0.160 |
| Context [Syn21] × Cell Line [Hsu..Cu..quinquefasciatus.] | 2.08 | 1.40 – 3.10 | **<0.001** |
| Context [BmLo] × Cell Line [Sf9..S..frugiperda.] | 0.71 | 0.48 – 1.06 | 0.093 |
| Context [Kozak] × Cell Line [Sf9..S..frugiperda.] | 0.92 | 0.62 – 1.37 | 0.689 |
| Context [Lep] × Cell Line [Sf9..S..frugiperda.] | 0.73 | 0.49 – 1.08 | 0.112 |
| TIS [Syn21] × Cell Line [Sf9..S..frugiperda.] | 2.17 | 1.46 – 3.22 | **<0.001** |
| TIS [BmLo] × Cell Line [U4.4..Ae..albopictus.] | 1.65 | 1.11 – 2.46 | **0.013** |
| TIS [Kozak] × Cell Line [U4.4..Ae..albopictus.] | 1.25 | 0.84 – 1.86 | 0.267 |
| TIS [Lep] × Cell Line [U4.4..Ae..albopictus.] | 1.23 | 0.83 – 1.83 | 0.304 |
| TIS [Syn21] × Cell Line [U4.4..Ae..albopictus.] | 1.62 | 1.09 – 2.40 | **0.017** |
| UTR [P10] × Cell Line [C6.36..Ae..albopictus.] | 0.47 | 0.32 – 0.71 | **<0.001** |
| UTR [SV40] × Cell Line [C6.36..Ae..albopictus.] | 1.72 | 1.15 – 2.55 | **0.007** |
| UTR [P10] × Cell Line [Hsu..Cu..quinquefasciatus.] | 0.73 | 0.49 – 1.08 | 0.117 |
| UTR [SV40] × Cell Line [Hsu..Cu..quinquefasciatus.] | 1.36 | 0.92 – 2.02 | 0.127 |
| UTR [P10] × Cell Line [Sf9..S..frugiperda.] | 5.43 | 3.66 – 8.07 | **<0.001** |
| UTR [SV40] × Cell Line [Sf9..S..frugiperda.] | 0.65 | 0.44 – 0.96 | **0.032** |
| UTR [P10] × Cell Line [U4.4..Ae..albopictus.] | 0.54 | 0.37 – 0.81 | **0.003** |
| UTR [SV40] × Cell Line [U4.4..Ae..albopictus.] | 1.11 | 0.75 – 1.65 | 0.596 |
| (TIS [BmLo] × UTR [P10]) × Cell Line [C6.36..Ae..albopictus.] | 0.95 | 0.54 – 1.66 | 0.850 |
| (TIS [Kozak] × UTR [P10]) × Cell Line [C6.36..Ae..albopictus.] | 0.45 | 0.25 – 0.78 | **0.005** |
| (TIS [Lep] × UTR [P10]) × Cell Line [C6.36..Ae..albopictus.] | 0.68 | 0.39 – 1.20 | 0.181 |
| (TIS [Syn21] × UTR [P10]) × Cell Line [C6.36..Ae..albopictus.] | 0.73 | 0.41 – 1.27 | 0.262 |
| (TIS [BmLo] × UTR [SV40]) × Cell Line [C6.36..Ae..albopictus.] | 1.65 | 0.94 – 2.90 | 0.078 |
| (TIS [Kozak] × UTR [SV40]) × Cell Line [C6.36..Ae..albopictus.] | 0.45 | 0.26 – 0.79 | **0.005** |
| (TIS [Lep] × UTR [SV40]) × Cell Line [C6.36..Ae..albopictus.] | 0.34 | 0.20 – 0.60 | **<0.001** |
| (TIS [Syn21] × UTR [SV40]) × Cell Line [C6.36..Ae..albopictus.] | 0.36 | 0.21 – 0.63 | **<0.001** |
| (TIS [BmLo] × UTR [P10]) × Cell Line [Hsu..Cu..quinquefasciatus.] | 0.71 | 0.41 – 1.24 | 0.230 |
| (TIS [Kozak] × UTR [P10]) × Cell Line [Hsu..Cu..quinquefasciatus.] | 1.34 | 0.76 – 2.34 | 0.309 |
| (TIS [Lep] × UTR [P10]) × Cell Line [Hsu..Cu..quinquefasciatus.] | 1.12 | 0.64 – 1.96 | 0.694 |
| (TIS [Syn21] × UTR [P10]) × Cell Line [Hsu..Cu..quinquefasciatus.] | 1.25 | 0.71 – 2.18 | 0.441 |
| (TIS [BmLo] × UTR [SV40]) × Cell Line [Hsu..Cu..quinquefasciatus.] | 0.86 | 0.49 – 1.50 | 0.587 |
| (TIS [Kozak] × UTR [SV40]) × Cell Line [Hsu..Cu..quinquefasciatus.] | 0.90 | 0.52 – 1.58 | 0.723 |
| (TIS [Lep] × UTR [SV40]) × Cell Line [Hsu..Cu..quinquefasciatus.] | 1.08 | 0.62 – 1.89 | 0.784 |
| (TIS [Syn21] × UTR [SV40]) × Cell Line [Hsu..Cu..quinquefasciatus.] | 0.70 | 0.40 – 1.22 | 0.207 |
| (TIS [BmLo] × UTR [P10]) × Cell Line [Sf9..S..frugiperda.] | 1.36 | 0.78 – 2.39 | 0.277 |
| (TIS [Kozak] × UTR [P10]) × Cell Line [Sf9..S..frugiperda.] | 1.37 | 0.78 – 2.40 | 0.270 |
| (TIS [Lep] × UTR [P10]) × Cell Line [Sf9..S..frugiperda.] | 0.85 | 0.48 – 1.48 | 0.558 |
| (TIS [Syn21] × UTR [P10]) × Cell Line [Sf9..S..frugiperda.] | 0.82 | 0.47 – 1.44 | 0.498 |
| (TIS [BmLo] × UTR [SV40]) × Cell Line [Sf9..S..frugiperda.] | 2.70 | 1.54 – 4.72 | **0.001** |
| (TIS [Kozak] × UTR [SV40]) × Cell Line [Sf9..S..frugiperda.] | 0.60 | 0.35 – 1.06 | 0.078 |
| (TIS [Lep] × UTR [SV40]) × Cell Line [Sf9..S..frugiperda.] | 0.71 | 0.40 – 1.24 | 0.223 |
| (TIS [Syn21] × UTR [SV40]) × Cell Line [Sf9..S..frugiperda.] | 0.49 | 0.28 – 0.85 | **0.011** |
| (TIS [BmLo] × UTR [P10]) × Cell Line [U4.4..Ae..albopictus.] | 0.56 | 0.32 – 0.99 | **0.045** |
| (TIS [Lep] × UTR [P10]) × Cell Line [U4.4..Ae..albopictus.] | 0.79 | 0.45 – 1.38 | 0.404 |
| (TIS [Syn21] × UTR [P10]) × Cell Line [U4.4..Ae..albopictus.] | 0.78 | 0.45 – 1.37 | 0.384 |
| (TIS [BmLo] × UTR [SV40]) × Cell Line [U4.4..Ae..albopictus.] | 1.38 | 0.79 – 2.42 | 0.260 |
| (TIS [Kozak] × UTR [SV40]) × Cell Line [U4.4..Ae..albopictus.] | 0.88 | 0.50 – 1.54 | 0.652 |
| (TIS [Lep] × UTR [SV40]) × Cell Line [U4.4..Ae..albopictus.] | 0.71 | 0.41 – 1.25 | 0.236 |
| (TIS [Syn21] × UTR [SV40]) × Cell Line [U4.4..Ae..albopictus.] | 0.63 | 0.36 – 1.10 | 0.101 |
| Observations | 592 | | |
| R^2^ Nagelkerke | 0.992 | | |

**Table S2. Pairwise contrasts table calculated from coefficients in full model (Table S1), back-calculated into a ratio difference on the original response scale.**

| Contrast | Mean Relative Expression | ±95% CI |
| --- | --- | --- |
| K10 / P10 | 0.20 | 0.19-0.22 |
| K10 / SV40 | 0.71 | 0.66-0.76 |
| P10 / SV40 | 3.47 | 3.22-3.73 |

| Contrast | Mean Relative Expression | ±95% CI |
| --- | --- | --- |
| BmHi / BmLo | 2.35 | 2.11-2.62 |
| BmHi / Kozak | 1.04 | 0.93-1.16 |
| BmHi / Lep | 0.92 | 0.83-1.03 |
| BmHi / Syn21 | 1.52 | 1.36-1.69 |
| BmLo / Kozak | 0.44 | 0.39-0.49 |
| BmLo / Lep | 0.39 | 0.35-0.44 |
| BmLo / Syn21 | 0.65 | 0.58-0.72 |
| Kozak / Lep | 0.89 | 0.79-0.99 |
| Kozak / Syn21 | 1.46 | 1.31-1.64 |
| Lep / Syn21 | 1.65 | 1.48-1.84 |

**Table S3. ANOVA table F values calculated from stepwise removal of coefficients from full model (Table S1).**

| Term | Df | F value | Pr(>F) |
| --- | --- | --- | --- |
| TIS | 4 | 36.43 | < 0.001 |
| UTR | 2 | 299.72 | < 0.001 |
| Residuals | 581 |  |  |
| Cell_Line | 4 | 594.57 | < 0.001 |
| TIS:UTR | 8 | 15.9 | < 0.001 |
| TIS:Cell_Line | 16 | 8.57 | < 0.001 |
| UTR:Cell_Line | 8 | 194.57 | < 0.001 |
| Residuals | 549 |  |  |
| TIS:UTR:Cell_Line | 31 | 4.2 | < 0.001 |
| Residuals | 518 |  |  |

##### Experiment Two

**Table S4. Fixed effects table for the generalized linear model (GLMM) fitted to the Luciferase Ratio detected in the engineered plasmids in experiment two– testing a range of promoters with a fixed combination of Contexts (TIS, 3’UTR) and two cell types**

|  | **Value** | | |
| --- | --- | --- | --- |
| *Predictors* | *Estimates* | *CI* | *p* |
| (Intercept) | 5.05 | 4.21 – 6.13 | **<0.001** |
| Promoter [OpIE2] | 0.10 | 0.07 – 0.13 | **<0.001** |
| Promoter [shortHsp83] | 0.08 | 0.06 – 0.10 | **<0.001** |
| Promoter [Hsp83] | 0.06 | 0.04 – 0.08 | **<0.001** |
| Promoter [CqPUb] | 0.00 | 0.00 – 0.00 | **<0.001** |
| Promoter [400AePUb] | 0.05 | 0.04 – 0.06 | **<0.001** |
| Promoter [800AePUb] | 0.47 | 0.36 – 0.62 | **<0.001** |
| Promoter [AePUb] | 2.19 | 1.68 – 2.87 | **<0.001** |
| Promoter [AlbPUb] | 1.49 | 1.18 – 1.88 | **0.001** |
| Context [Kozak_SV40] | 3.76 | 2.97 – 4.72 | **<0.001** |
| Context [Lep_P10] | 12.98 | 9.94 – 16.96 | **<0.001** |
| Rep [MA34] | 1.00 | 0.76 – 1.30 | 0.976 |
| Rep [MA37] | 0.54 | 0.42 – 0.71 | **<0.001** |
| Cell Line [U4] | 0.73 | 0.56 – 0.96 | **0.023** |
| Promoter [OpIE2] × Context [Kozak_SV40] | 0.47 | 0.33 – 0.68 | **<0.001** |
| Promoter [shortHsp83] × Context [Kozak_SV40] | 1.17 | 0.83 – 1.67 | 0.373 |
| Promoter [Hsp83] × Context [Kozak_SV40] | 1.59 | 1.12 – 2.26 | **0.010** |
| Promoter [CqPUb] × Context [Kozak_SV40] | 2.32 | 1.55 – 3.45 | **<0.001** |
| Promoter [400AePUb] × Context [Kozak_SV40] | 0.82 | 0.57 – 1.16 | 0.262 |
| Promoter [800AePUb] × Context [Kozak_SV40] | 0.85 | 0.60 – 1.22 | 0.378 |
| Promoter [AePUb] × Context [Kozak_SV40] | 1.49 | 1.05 – 2.13 | **0.026** |
| Promoter [OpIE2] × Context [Lep_P10] | 1.55 | 1.06 – 2.26 | **0.023** |
| Promoter [shortHsp83] × Context [Lep_P10] | 1.12 | 0.77 – 1.64 | 0.548 |
| Promoter [Hsp83] × Context [Lep_P10] | 1.30 | 0.89 – 1.89 | 0.176 |
| Promoter [CqPUb] × Context [Lep_P10] | 3.79 | 2.47 – 5.76 | **<0.001** |
| Promoter [400AePUb] × Context [Lep_P10] | 0.55 | 0.38 – 0.81 | **0.002** |
| Promoter [800AePUb] × Context [Lep_P10] | 0.56 | 0.39 – 0.82 | **0.003** |
| Promoter [AePUb] × Context [Lep_P10] | 0.68 | 0.47 – 0.99 | **0.044** |
| Promoter [OpIE2] × Rep [MA34] | 1.39 | 0.95 – 2.03 | 0.087 |
| Promoter [shortHsp83] × Rep [MA34] | 0.75 | 0.52 – 1.10 | 0.138 |
| Promoter [Hsp83] × Rep [MA34] | 0.72 | 0.50 – 1.06 | 0.093 |
| Promoter [CqPUb] × Rep [MA34] | 0.36 | 0.23 – 0.54 | **<0.001** |
| Promoter [400AePUb] × Rep [MA34] | 0.60 | 0.41 – 0.88 | **0.008** |
| Promoter [800AePUb] × Rep [MA34] | 0.48 | 0.33 – 0.69 | **<0.001** |
| Promoter [AePUb] × Rep [MA34] | 0.88 | 0.60 – 1.28 | 0.503 |
| Promoter [AlbPUb] × Rep [MA34] | 0.75 | 0.54 – 1.04 | 0.080 |
| Promoter [OpIE2] × Rep [MA37] | 0.98 | 0.67 – 1.43 | 0.906 |
| Promoter [shortHsp83] × Rep [MA37] | 2.47 | 1.69 – 3.60 | **<0.001** |
| Promoter [Hsp83] × Rep [MA37] | 2.42 | 1.66 – 3.53 | **<0.001** |
| Promoter [CqPUb] × Rep [MA37] | 1.69 | 1.11 – 2.57 | **0.014** |
| Promoter [400AePUb] × Rep [MA37] | 1.05 | 0.72 – 1.53 | 0.796 |
| Promoter [800AePUb] × Rep [MA37] | 0.84 | 0.58 – 1.23 | 0.367 |
| Promoter [AePUb] × Rep [MA37] | 1.31 | 0.90 – 1.92 | 0.157 |
| Promoter [AlbPUb] × Rep [MA37] | 1.64 | 1.19 – 2.28 | **0.003** |
| Context [Kozak_SV40] × Rep [MA34] | 0.65 | 0.47 – 0.91 | **0.011** |
| Context [Lep_P10] × Rep [MA34] | 0.78 | 0.54 – 1.14 | 0.206 |
| Context [Kozak_SV40] × Rep [MA37] | 1.51 | 1.09 – 2.09 | **0.014** |
| Context [Lep_P10] × Rep [MA37] | 2.54 | 1.74 – 3.70 | **<0.001** |
| Promoter [OpIE2] × Cell Line [U4] | 0.98 | 0.67 – 1.44 | 0.935 |
| Promoter [shortHsp83] × Cell Line [U4] | 0.07 | 0.05 – 0.10 | **<0.001** |
| Promoter [Hsp83] × Cell Line [U4] | 0.10 | 0.07 – 0.15 | **<0.001** |
| Promoter [CqPUb] × Cell Line [U4] | 0.36 | 0.23 – 0.55 | **<0.001** |
| Promoter [400AePUb] × Cell Line [U4] | 0.07 | 0.05 – 0.10 | **<0.001** |
| Promoter [800AePUb] × Cell Line [U4] | 0.03 | 0.02 – 0.05 | **<0.001** |
| Promoter [AePUb] × Cell Line [U4] | 0.15 | 0.10 – 0.22 | **<0.001** |
| Context [Kozak_SV40] × Cell Line [U4] | 0.61 | 0.44 – 0.84 | **0.003** |
| Context [Lep_P10] × Cell Line [U4] | 0.40 | 0.27 – 0.58 | **<0.001** |
| Rep [MA34] × Cell Line [U4] | 1.60 | 1.10 – 2.34 | **0.014** |
| Rep [MA37] × Cell Line [U4] | 1.96 | 1.34 – 2.85 | **<0.001** |
| (Promoter [OpIE2] × Context [Kozak_SV40]) × Rep [MA34] | 0.93 | 0.56 – 1.53 | 0.771 |
| (Promoter [shortHsp83] × Context [Kozak_SV40]) × Rep [MA34] | 1.38 | 0.83 – 2.27 | 0.211 |
| (Promoter [Hsp83] × Context [Kozak_SV40]) × Rep [MA34] | 1.18 | 0.72 – 1.94 | 0.517 |
| (Promoter [CqPUb] × Context [Kozak_SV40]) × Rep [MA34] | 2.11 | 1.24 – 3.61 | **0.006** |
| (Promoter [400AePUb] × Context [Kozak_SV40]) × Rep [MA34] | 1.22 | 0.74 – 2.01 | 0.437 |
| (Promoter [800AePUb] × Context [Kozak_SV40]) × Rep [MA34] | 1.66 | 1.00 – 2.73 | **0.048** |
| (Promoter [AePUb] × Context [Kozak_SV40]) × Rep [MA34] | 0.83 | 0.51 – 1.38 | 0.478 |
| (Promoter [OpIE2] × Context [Lep_P10]) × Rep [MA34] | 0.78 | 0.46 – 1.33 | 0.362 |
| (Promoter [shortHsp83] × Context [Lep_P10]) × Rep [MA34] | 1.09 | 0.64 – 1.86 | 0.744 |
| (Promoter [Hsp83] × Context [Lep_P10]) × Rep [MA34] | 0.91 | 0.53 – 1.55 | 0.729 |
| (Promoter [CqPUb] × Context [Lep_P10]) × Rep [MA34] | 1.88 | 1.07 – 3.32 | **0.029** |
| (Promoter [400AePUb] × Context [Lep_P10]) × Rep [MA34] | 1.14 | 0.67 – 1.94 | 0.632 |
| (Promoter [800AePUb] × Context [Lep_P10]) × Rep [MA34] | 1.46 | 0.86 – 2.50 | 0.161 |
| (Promoter [AePUb] × Context [Lep_P10]) × Rep [MA34] | 0.92 | 0.54 – 1.56 | 0.748 |
| (Promoter [OpIE2] × Context [Kozak_SV40]) × Rep [MA37] | 1.21 | 0.73 – 1.99 | 0.459 |
| (Promoter [shortHsp83] × Context [Kozak_SV40]) × Rep [MA37] | 0.84 | 0.51 – 1.38 | 0.493 |
| (Promoter [Hsp83] × Context [Kozak_SV40]) × Rep [MA37] | 0.73 | 0.44 – 1.20 | 0.211 |
| (Promoter [CqPUb] × Context [Kozak_SV40]) × Rep [MA37] | 0.72 | 0.42 – 1.23 | 0.223 |
| (Promoter [400AePUb] × Context [Kozak_SV40]) × Rep [MA37] | 0.96 | 0.59 – 1.59 | 0.888 |
| (Promoter [800AePUb] × Context [Kozak_SV40]) × Rep [MA37] | 1.12 | 0.68 – 1.85 | 0.655 |
| (Promoter [AePUb] × Context [Kozak_SV40]) × Rep [MA37] | 0.34 | 0.21 – 0.56 | **<0.001** |
| (Promoter [OpIE2] × Context [Lep_P10]) × Rep [MA37] | 0.65 | 0.38 – 1.11 | 0.113 |
| (Promoter [shortHsp83] × Context [Lep_P10]) × Rep [MA37] | 0.38 | 0.22 – 0.65 | **<0.001** |
| (Promoter [Hsp83] × Context [Lep_P10]) × Rep [MA37] | 0.37 | 0.22 – 0.63 | **<0.001** |
| (Promoter [CqPUb] × Context [Lep_P10]) × Rep [MA37] | 0.49 | 0.28 – 0.86 | **0.012** |
| (Promoter [400AePUb] × Context [Lep_P10]) × Rep [MA37] | 0.48 | 0.28 – 0.81 | **0.006** |
| (Promoter [800AePUb] × Context [Lep_P10]) × Rep [MA37] | 0.62 | 0.36 – 1.06 | 0.078 |
| (Promoter [AePUb] × Context [Lep_P10]) × Rep [MA37] | 0.35 | 0.21 – 0.60 | **<0.001** |
| (Promoter [OpIE2] × Context [Kozak_SV40]) × Cell Line [U4] | 1.05 | 0.64 – 1.73 | 0.856 |
| (Promoter [shortHsp83] × Context [Kozak_SV40]) × Cell Line [U4] | 1.74 | 1.06 – 2.86 | **0.030** |
| (Promoter [Hsp83] × Context [Kozak_SV40]) × Cell Line [U4] | 1.02 | 0.62 – 1.67 | 0.951 |
| (Promoter [CqPUb] × Context [Kozak_SV40]) × Cell Line [U4] | 0.49 | 0.29 – 0.84 | **0.010** |
| (Promoter [400AePUb] × Context [Kozak_SV40]) × Cell Line [U4] | 1.27 | 0.77 – 2.09 | 0.351 |
| (Promoter [800AePUb] × Context [Kozak_SV40]) × Cell Line [U4] | 1.37 | 0.83 – 2.25 | 0.222 |
| (Promoter [AePUb] × Context [Kozak_SV40]) × Cell Line [U4] | 0.95 | 0.57 – 1.56 | 0.828 |
| (Promoter [OpIE2] × Context [Lep_P10]) × Cell Line [U4] | 0.94 | 0.55 – 1.60 | 0.816 |
| (Promoter [shortHsp83] × Context [Lep_P10]) × Cell Line [U4] | 3.19 | 1.87 – 5.43 | **<0.001** |
| (Promoter [Hsp83] × Context [Lep_P10]) × Cell Line [U4] | 2.52 | 1.48 – 4.29 | **0.001** |
| (Promoter [CqPUb] × Context [Lep_P10]) × Cell Line [U4] | 1.68 | 0.95 – 2.97 | 0.076 |
| (Promoter [400AePUb] × Context [Lep_P10]) × Cell Line [U4] | 2.67 | 1.56 – 4.55 | **<0.001** |
| (Promoter [800AePUb] × Context [Lep_P10]) × Cell Line [U4] | 6.72 | 3.94 – 11.46 | **<0.001** |
| (Promoter [AePUb] × Context [Lep_P10]) × Cell Line [U4] | 4.10 | 2.40 – 6.98 | **<0.001** |
| (Promoter [OpIE2] × Rep [MA34]) × Cell Line [U4] | 0.47 | 0.28 – 0.81 | **0.006** |
| (Promoter [shortHsp83] × Rep [MA34]) × Cell Line [U4] | 2.58 | 1.51 – 4.40 | **0.001** |
| (Promoter [Hsp83] × Rep [MA34]) × Cell Line [U4] | 2.01 | 1.18 – 3.42 | **0.010** |
| (Promoter [CqPUb] × Rep [MA34]) × Cell Line [U4] | 0.81 | 0.46 – 1.44 | 0.478 |
| (Promoter [400AePUb] × Rep [MA34]) × Cell Line [U4] | 0.71 | 0.41 – 1.21 | 0.203 |
| (Promoter [800AePUb] × Rep [MA34]) × Cell Line [U4] | 1.53 | 0.90 – 2.61 | 0.117 |
| (Promoter [AePUb] × Rep [MA34]) × Cell Line [U4] | 0.56 | 0.33 – 0.96 | **0.033** |
| (Promoter [OpIE2] × Rep [MA37]) × Cell Line [U4] | 1.02 | 0.60 – 1.75 | 0.929 |
| (Promoter [shortHsp83] × Rep [MA37]) × Cell Line [U4] | 0.99 | 0.58 – 1.68 | 0.960 |
| (Promoter [Hsp83] × Rep [MA37]) × Cell Line [U4] | 0.85 | 0.50 – 1.45 | 0.543 |
| (Promoter [CqPUb] × Rep [MA37]) × Cell Line [U4] | 0.25 | 0.14 – 0.45 | **<0.001** |
| (Promoter [400AePUb] × Rep [MA37]) × Cell Line [U4] | 0.93 | 0.54 – 1.58 | 0.785 |
| (Promoter [800AePUb] × Rep [MA37]) × Cell Line [U4] | 1.42 | 0.83 – 2.42 | 0.196 |
| (Promoter [AePUb] × Rep [MA37]) × Cell Line [U4] | 0.87 | 0.51 – 1.48 | 0.601 |
| (Context [Kozak_SV40] × Rep [MA34]) × Cell Line [U4] | 1.47 | 0.92 – 2.33 | 0.104 |
| (Context [Lep_P10] × Rep [MA34]) × Cell Line [U4] | 1.21 | 0.71 – 2.06 | 0.487 |
| (Context [Kozak_SV40] × Rep [MA37]) × Cell Line [U4] | 1.03 | 0.65 – 1.64 | 0.888 |
| (Context [Lep_P10] × Rep [MA37]) × Cell Line [U4] | 0.52 | 0.31 – 0.89 | **0.017** |
| (Promoter [OpIE2] × Context [Kozak_SV40] × Rep [MA34]) × Cell Line [U4] | 1.36 | 0.67 – 2.75 | 0.394 |
| (Promoter [shortHsp83] × Context [Kozak_SV40] × Rep [MA34]) × Cell Line [U4] | 0.77 | 0.38 – 1.55 | 0.457 |
| (Promoter [Hsp83] × Context [Kozak_SV40] × Rep [MA34]) × Cell Line [U4] | 1.08 | 0.54 – 2.20 | 0.822 |
| (Promoter [CqPUb] × Context [Kozak_SV40] × Rep [MA34]) × Cell Line [U4] | 1.23 | 0.59 – 2.57 | 0.576 |
| (Promoter [400AePUb] × Context [Kozak_SV40] × Rep [MA34]) × Cell Line [U4] | 1.36 | 0.67 – 2.75 | 0.395 |
| (Promoter [800AePUb] × Context [Kozak_SV40] × Rep [MA34]) × Cell Line [U4] | 0.49 | 0.24 – 1.00 | 0.051 |
| (Promoter [AePUb] × Context [Kozak_SV40] × Rep [MA34]) × Cell Line [U4] | 1.22 | 0.60 – 2.46 | 0.586 |
| (Promoter [OpIE2] × Context [Lep_P10] × Rep [MA34]) × Cell Line [U4] | 1.51 | 0.71 – 3.21 | 0.284 |
| (Promoter [shortHsp83] × Context [Lep_P10] × Rep [MA34]) × Cell Line [U4] | 0.58 | 0.27 – 1.22 | 0.151 |
| (Promoter [Hsp83] × Context [Lep_P10] × Rep [MA34]) × Cell Line [U4] | 0.74 | 0.35 – 1.57 | 0.433 |
| (Promoter [CqPUb] × Context [Lep_P10] × Rep [MA34]) × Cell Line [U4] | 1.26 | 0.58 – 2.77 | 0.557 |
| (Promoter [400AePUb] × Context [Lep_P10] × Rep [MA34]) × Cell Line [U4] | 1.07 | 0.50 – 2.28 | 0.857 |
| (Promoter [800AePUb] × Context [Lep_P10] × Rep [MA34]) × Cell Line [U4] | 0.37 | 0.17 – 0.78 | **0.009** |
| (Promoter [AePUb] × Context [Lep_P10] × Rep [MA34]) × Cell Line [U4] | 1.45 | 0.68 – 3.08 | 0.334 |
| (Promoter [OpIE2] × Context [Kozak_SV40] × Rep [MA37]) × Cell Line [U4] | 0.85 | 0.42 – 1.72 | 0.647 |
| (Promoter [shortHsp83] × Context [Kozak_SV40] × Rep [MA37]) × Cell Line [U4] | 0.82 | 0.41 – 1.67 | 0.587 |
| (Promoter [Hsp83] × Context [Kozak_SV40] × Rep [MA37]) × Cell Line [U4] | 1.41 | 0.70 – 2.86 | 0.336 |
| (Promoter [CqPUb] × Context [Kozak_SV40] × Rep [MA37]) × Cell Line [U4] | 3.27 | 1.57 – 6.80 | **0.002** |
| (Promoter [400AePUb] × Context [Kozak_SV40] × Rep [MA37]) × Cell Line [U4] | 1.08 | 0.53 – 2.18 | 0.838 |
| (Promoter [800AePUb] × Context [Kozak_SV40] × Rep [MA37]) × Cell Line [U4] | 0.96 | 0.48 – 1.95 | 0.919 |
| (Promoter [AePUb] × Context [Kozak_SV40] × Rep [MA37]) × Cell Line [U4] | 2.71 | 1.34 – 5.49 | **0.006** |
| (Promoter [OpIE2] × Context [Lep_P10] × Rep [MA37]) × Cell Line [U4] | 1.29 | 0.61 – 2.74 | 0.512 |
| (Promoter [shortHsp83] × Context [Lep_P10] × Rep [MA37]) × Cell Line [U4] | 1.37 | 0.64 – 2.92 | 0.412 |
| (Promoter [Hsp83] × Context [Lep_P10] × Rep [MA37]) × Cell Line [U4] | 1.70 | 0.80 – 3.61 | 0.170 |
| (Promoter [CqPUb] × Context [Lep_P10] × Rep [MA37]) × Cell Line [U4] | 3.17 | 1.45 – 6.91 | **0.004** |
| (Promoter [400AePUb] × Context [Lep_P10] × Rep [MA37]) × Cell Line [U4] | 1.59 | 0.75 – 3.38 | 0.228 |
| (Promoter [800AePUb] × Context [Lep_P10] × Rep [MA37]) × Cell Line [U4] | 0.81 | 0.38 – 1.72 | 0.584 |
| (Promoter [AePUb] × Context [Lep_P10] × Rep [MA37]) × Cell Line [U4] | 2.86 | 1.34 – 6.07 | **0.006** |
| Observations | 1218 | | |
| R^2^ Nagelkerke | 0.999 | | |

**Table S5. ANOVA table F values calculated from stepwise removal of coefficients from full model (Table S4).**

| Term | Df | F value | Pr(>F) |
| --- | --- | --- | --- |
| Promoter | 8 | 354.91 | < 0.001 |
| Context | 2 | 509.96 | < 0.001 |
| Residuals | 1207 |  |  |
| Promoter: Context | 14 | 7.255 | < 0.001 |
| Residuals | 1193 |  |  |

### Methods

All cell lines were maintained at 28˚C, without CO2 or humidity control. Aag2, U4.4 and C6/36 cells were maintained in L-15 (Thermo Fisher Scientific, Waltham, MA, US) supplemented with 10% FBS (Labtech, Lewes, UK), 1%Pen/Strep (Thermo Fisher Scientific, Waltham, MA, US) and 10% Tryptose Phosphate Broth (Thermo Fisher Scientific, Waltham, MA, US). Hsu cells were maintained in Schneider’s Drosophila Medium (Lonza, Basel, Switzerland) supplemented with 10% FBS (Labtech, Lewes, UK) and 1% Pen/Strep (Thermo Fisher Scientific, Waltham, MA, US). Sf9 cells were maintained in Insect Xpress (Lonza, Basel, Switzerland) supplemented with 10% FBS (Labtech, Lewes, UK) and 1% Pen/Strep (Thermo Fisher Scientific, Waltham, MA, US).

| **Table S6. Transfection amounts per cell line** |  |  |  |  |  |
| --- | --- | --- | --- | --- | --- |
|  | Aag2 | C6.36 | Hsu | Sf9 | U4.4 |
| FF plasmid (ng/well) | 1 | 1 | 1 | 5 | 1 |
| RL plasmid (ng/well) | 50 | 5 | 5 | 1 | 5 |
| TransIT Pro reagent (μl/well) | 0.2 | 0.2 | 0.2 | 0.2 | 0.2 |
| TransIT Boost reagent (μl/well) | 0.1 | 0.1 | 0.1 | 0.1 | 0.1 |

##### Dual luciferase assay

Within an experiment, different cell lysate volumes (usually two) were screened for each cell line; and were used to select a lysate volume for the rest of the samples in that cell line. Each single luciferase control is represented, as is a double luciferase sample that is expected to have high expression of firefly luciferase (FF). The same two repeats are used at each lysate volume, but the FF plasmid in the double positive sample (“FF and RL”) is not the same as that in the single positive (“FF only”) sample.

In Figure S1 the FF and RL measurements of each sample are indicated as different colours with the transfection condition (combination of plasmids) represented on the x-axis and arbitrary light units (ALU) on the y-axis. The two lysate volumes, 1µl and 4µl are represented on independent graphs with the cell line indicated above (Aag2, in this case). Although the specific FF quenching threshold (106 ALU) is indicated, an estimated RL background threshold must be used as there is not enough data to calculate the 99.9% confidence interval it is based upon.

In optimisation experiment 2 (Figure S1), both lysate volumes show the desired pattern of results: each luciferase activity present above background threshold only where the corresponding luciferase plasmid was present in the transfection and FF activity present below the FF quenching threshold (106 ALU). It was decided to proceed with 1µl lysate volume for the rest of the experiment as RL measurements are higher without corresponding FF measurements becoming close to the quenching threshold (there may be experimental samples with greater FF activity than in the samples screened here). The volumes used are given in Table S7.

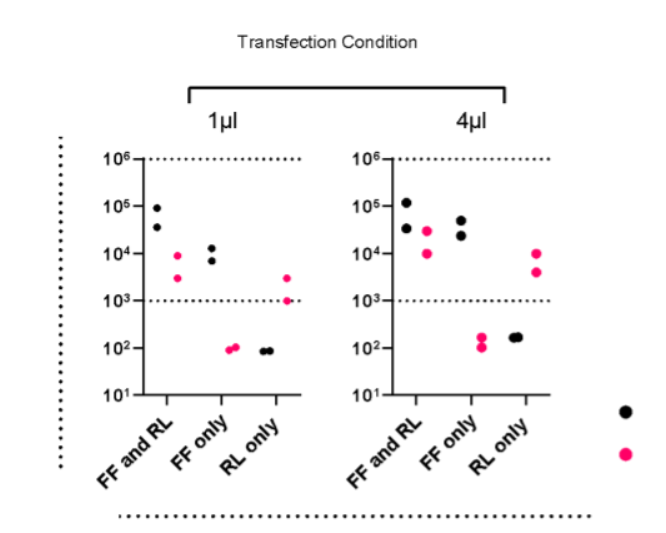

**Figure S1. Indicative lysate volume. Graphs of lysate controls for optimisation experiments. Samples of control transfections were processed by dual luciferase assay at two volumes (1 and 4 µl). Each represented on a separate panel, with Arbitrary Light Units (ALU) on the y-axis.**

**Table S7. Indicative lysate volume for experiments**

|  | Lysate volume (µl) | | | | |
| --- | --- | --- | --- | --- | --- |
|  | Aag2 | C6.36 | Hsu | Sf9 | U4.4 |
| Optimisation experiment 1 | 1 | - | - | - | 1 |
| Optimisation experiment 2 | 4 | - | - | - | - |
| Experiment | 1 | 1 | 1 | 1 | 7 |

##### TIS sequences

**Table S8. Five translation initiation sequences used in this project.**

| Name | Sequence |
| --- | --- |
| Kozak (Koz) | GCCGCCACC**ATG**G |
| Lepidopteran (Lep) | AACCAACAAC**ATG**G |
| *B. mori* High (BmHi) | AAAAATCAAA**ATG**G |
| *B. mori* Low (BmLo) | CCGCCGGCGT**ATG**G |
| Syn21 (Syn21) | AACTTAAAAAAAAAAATCAAA**ATG**G |

**Table S9. Brief descriptors and Lab plasmid registry names**

| **Name** | **AGG number** |
| --- | --- |
| HR5-IE1-Koz-FF-SV40 | AGG1186 |
| HR5-IE1-Lep-FF-SV40 | AGG1187 |
| HR5-IE1-BmHi-FF-SV40 | AGG1188 |
| HR5-IE1-BmLo-FF-SV40 | AGG1189 |
| HR5-IE1-Koz-FF-K10 | AGG1191 |
| HR5-IE1-Lep-FF-K10 | AGG1192 |
| HR5-IE1-BmHi-FF-K10 | AGG1193 |
| HR5-IE1-BmLo-FF-K10 | AGG1194 |
| HR5-IE1-Koz-FF-P10 | AGG1196 |
| HR5-IE1-Lep-FF-P10 | AGG1197 |
| HR5-IE1-BmHi-FF-P10 | AGG1198 |
| HR5-IE1-BmLo-FF-P10 | AGG1199 |
| AePUB-Kozak-FF-SV40 | AGG1330 |
| AePUB-Lep-FF-P10 | AGG1331 |
| AePUB-BmLo-FF-K10 | AGG1332 |
| 400AePUB-Kozak-FF-SV40 | AGG1333 |
| 400AePUB-Lep-FF-P10 | AGG1334 |
| 400AePUB-B.mori_low-FF-K10 | AGG1335 |
| 800AePUB-Kozak-FF-SV40 | AGG1336 |
| 800AePUB-Lep-FF-P10 | AGG1337 |
| 800AePUB-BmLo-FF-K10 | AGG1338 |
| AlbPUb-Kozak-FF-SV40 | AGG1345 |
| Hsp83-Kozak-FF-SV40 | AGG1390 |
| Hsp83-Lep-FF-P10 | AGG1391 |
| Hsp83-BmLo-FF-K10 | AGG1392 |
| sHsp83-Kozak-FF-SV40 | AGG1393 |
| sHsp83-Lep-FF-P10 | AGG1394 |
| shortthsp83-BmLo-FF-K10 | AGG1395 |
| OPIE2-Kozak-FF-SV40 | AGG1396 |
| OPIE2-Lep-FF-P10 | AGG1397 |
| OPIE2-BmLo-FF-K10 | AGG1398 |
| CQ PUb - Kozak - SV40 | AGG1416 |
| CQ PUb- Lep-FF - P10 | AGG1417 |
| CQ PUb – BmLo-FF-K10 | AGG1418 |
